# Night-time neuronal activation of Cluster N in a North American songbird

**DOI:** 10.1101/2023.06.07.544090

**Authors:** Jennifer Rudolf, Natalie Philipello, Tamara Fleihan, J. David Dickman, Kira E. Delmore

## Abstract

Night-migrating songbirds utilize the Earth’s magnetic field to help navigate to and from their breeding sites each year. A region of the avian forebrain called Cluster N has been shown to be activated during night migratory behavior and it has been implicated in processing geomagnetic information. Previous studies with night-migratory European songbirds have shown that neuronal activity at Cluster N is higher at night than during the day. Comparable work in North American migrants has only been performed in one species of swallows, so extension of examination for Cluster N in other migratory birds is needed. In addition, it is unclear if Cluster N activation is lateralized and the full extent of its boundaries in the forebrain have yet to be described. We used sensory-driven gene expression based on ZENK and the Swainson’s thrush, a night-migratory North American songbird, to fill these knowledge gaps. We found elevated levels of gene expression in night-vs. day-active thrushes and no evidence for lateralization in this region. We further examined the anatomical extent of neural activation in the forebrain using 3D reconstruction topology. Our findings demonstrate that Swainson’s thrushes possess an extensive bilateral night-activated Cluster N region in the forebrain similar to other European avian species, suggesting that Cluster N is highly conserved in nocturnal migrants.

## Introduction

Migratory birds can travel thousands of miles between their breeding sites and wintering sites twice each year. How migratory birds accomplish these long-distance navigations remains unknown, but it appears that the brain uses several cues including visual, olfactory, and a sense of the Earth’s magnetic field. Many of these navigating birds travel at night, and a number of investigations have established a behavioral pattern of migratory restlessness where animals orient in the direction of desired travel during the night hours but not during the day [3,37]. One region of the avian brain that has been implicated in migratory behavior is termed Cluster N [22,31,46]. Cluster N is a broad region in the dorsal forebrain of birds that has largely been identified through increased levels of neural activity during migratory behaviors or magnetic field manipulation [6,26,31]. Although never fully characterized, Cluster N has been localized anatomically to be contained in the avian visual Wulst and includes portions of the hyperpallium, dorsal mesopallium, and hippocampal formation. Cluster N receives a direct projection from the visual related dorsal geniculate nucleus of the thalamus and at least some Cluster N neurons have been shown to be di-synaptically connected to the retina through the thalamofugal pathway [22,29]. Previous studies in migrating birds have shown elevated neuronal activity levels in Cluster N at night but not during the day [6,46]. Comparable increases in neural activity were not noted in non-migratory birds [26,31,46], or in birds not exhibiting migratory restlessness behavior in the dark [6]. Chemical inactivation of Cluster N in European robins disrupted magnetic orientation behavior and decreased Cluster N neural activity was described when birds eyes were covered during migratory behavior [26,31,45].

To the best of our knowledge, existing work on Cluster N has been focused on European and holarctic species, including the garden warbler (*Sylvia borin*), European robin (*Erithacus rubecula;)*, meadow pipit (*Anthus pratensis*) and the northern wheatear (*Oenanthe oenanthe*) [17,31,46], except for one study in North American sparrows [6]. The reliability of different compass mechanisms can change depending on location, direction of travel, and season suggesting additional work outside Europe is needed [32]. Additional gaps in our knowledge of Cluster N also exist. For example, it is unclear how the region is activated only during migratory season, only at night, and how magnetic field information is processed by Cluster N neurons for navigation behavior. It has been suggested that Cluster N may only be active in the right hemisphere of migrating birds [26,36,42,43], but other studies have not supported unilateral hemispheric activation [18,20,21,41]. In addition, the neuroanatomical and physiological properties of Cluster N cells in the avian forebrain as characterized by migratory activation has yet to be provided. Work presented here will extend these knowledge gaps by examining Cluster N in a North American migratory species, the Swainson’s thrush (*Catharus ustulatus*).

The Swainson’s thrush is a night-migrating songbird that breeds in North America and winters in Mexico, Central America, and South America. There is strong evidence Swainson’s thrushes use magnetic cues to orient on migration, with large scale tracking of free-flying birds showing that birds exposed to altered magnetic fields orient in the wrong direction. Birds were able to re-orient on subsequent nights using twilight cues and the true magnetic field [8,9]. Here, we used sensory-driven gene expression and comparisons between day-and night-active birds to define Cluster N in Swainson’s thrushes during migratory behavior. We had three specific aims, to (1) test for increased neuronal activity in the region described as Cluster N in previous studies, (2) test for lateralization in this region, and (3) generate a 3D representation of Cluster N showing its size and location in the forebrain. We compared levels of neural activation in the forebrain during night and day activity during migration season, examined differences in hemispheric expression, and generated a 3D representation of Cluster N.

## Methods

All protocols outlined in this study were approved by the Institutional Animal Care and Use Committee at Texas A&M (IACUC 2019-0066) and permits were obtained from Environment and Climate Change Canada (SC-BC-2020-0016), the U.S. Fish and Wildlife Service (MB49986D-0) and Texas Parks and Wildlife Commission (SPR-0419-067).

### Experimental Procedures

We captured Swainson’s thrushes at the end of the breeding season (August 2020) in Kamloops, British Columbia, and transferred them to an indoor vivarium in College Station, Texas. We used data from birds fitted with light-level geolocators from the same sites in previous years to mimic natural conditions for captive birds, bringing them from 13 hours of daylight upon arrival in College Station to 11.5 hours in early October (Oct 8, ¼ hour reductions in daylight every 5 days starting Sep 10). We used infrared cameras to monitor birds at night to ensure they all exhibited nocturnal behavior by early October.

Experiments took place from October 13-17, 2020. We monitored the birds for 90 minutes before sacrifice, noting their behavior every five minutes. Birds were randomly placed into day- or night-active groups. For night experiments, only birds that exhibited *zugunruhe* (or migratory restlessness) for at least 60 minutes (occurred 120-140 minutes after lights out) were utilized for neural activity gene expression experiments. *Zugunruhe* was defined as movement between perches, hopping and flapping wings [2]. Other behaviors exhibited by both day- and night-active birds that would not have been considered *zugunruhe* included preening, jumping between perches, head movements, still but awake on perch, feeding. Four night- and two day-active birds were used to examine ZENK expression. Day active birds were euthanized at midday and were not exhibiting *zugunruhe* (11AM - 1 PM). We deeply anesthetized the birds with sodium pentobarbital and transcardially perfused with saline and 4% paraformaldehyde. Birds were decapitated, brains extracted, hemispheres separated and immersed them in 4% PFA for 12 hours. We then transferred the hemispheres to 30% sucrose for 24 hours before serially sectioning and storing in 0.1 M PBS at 4 degrees C. We sectioned the brains sagittally at 40 μm.

### Immunohistochemistry

We used ZENK to quantify neuronal activity induced gene transcription. ZENK is a transcription factor whose expression increases rapidly when cells are stimulated (i.e., when synaptic activity is occurring in the brain) and is commonly used in birds to analyze brain activity [27,28]. We used the monoclonal antibody mouse mAb (7B7-A3) validated in three avian species: the rock pigeon, zebra finch and domestic chicken [34]. We incubated free floating sections of the brain in H2O2 (1:50) for 30 minutes then blocked it with 10% Normal Horse Serum (Vector Laboratories, Burlingame, CA, USA) for one hour. We incubated the sections in our primary antibody, ZENK mouse mAb (1:5000 in TBS-T), for 24 hours, secondary antibody (Horse Anti-Mouse IgG, 1:500 in TBS-T; Vector Laboratories, Burlingame, CA, USA), for 1 hour, avidin-biotin horseradish peroxidase-complex (1:200 in TBS, Vectastain Elite Kit, Vector Laboratories, Burlingame, CA USA) for one hour, then reacted for visualization of peroxidase with nickel-cobalt intensified diaminobenzidine for 3 - 8 minutes (DAB Peroxidase substrate kit SK-4100, Vector Laboratories, Burlingame, CA, USA). Brain slices were washed between each incubation step using TBS (3 × 10 minutes) and mounted on slides before being stained with Neutral red.

In alternate sections for 2 birds, Anti-Glutamate Receptor 1 (GluR1) was used to aid in establishing the neuroanatomical subdivisions of the brain [39]. We followed the same procedures described above for ZENK, replacing the Normal Horse Serum for 10% Normal Goat Serum, and using Glu-R1 (Millipore Cat# AB1504, 1:250; Vector Laboratories, Burlingame, CA, USA) for the primary and Anti-glutamate receptor 1 (rabbit polyclonal) Goat Anti-rabbit HRP (Novus Biologicals Cat# NB7160, 1:250; Vector Laboratories, Burlingame, CA, USA) for the secondary antibodies, respectively.

### Quantifying immunolabeled cells

We scanned stained slides using an Aperio Scanscope3 system (Aperio, Vista, CA) at 20× magnification and used Imagescope software to annotate two regions on each brain section: (1) the area expected to include cluster N (following previous studies and our own ZENK and GluR1 staining, Fig 1E) and (2) an area anterior to this region that was not expected to include cluster N (Fig 1F). All annotations were done by the same individual (NP) and naïve to the experimental conditions. We used an Aperio algorithm to identify cells with low, medium or high expression and summarized these data for each annotated region as the proportion of cells that exhibited medium and high expression (number medium + number high / total expressed cells). Finally, we subtracted the anterior proportion from the cluster N proportion to control for potential differences in background staining on each slide. The same approach was used in previous studies of cluster N [26,31,45]. We summarized cells using this method for both the left and right hemisphere of each bird. We used a linear mixed effect model to analyze these data, with condition (day-vs. night-active) as the predictor variable and bird as a random variable (accounting for the fact measures for both the left and right hemisphere were included). A one sample t-test was used to test for lateralization, calculating differences between the left and right hemisphere in each bird and comparing these values to an expected difference of zero.

**Fig 1.**
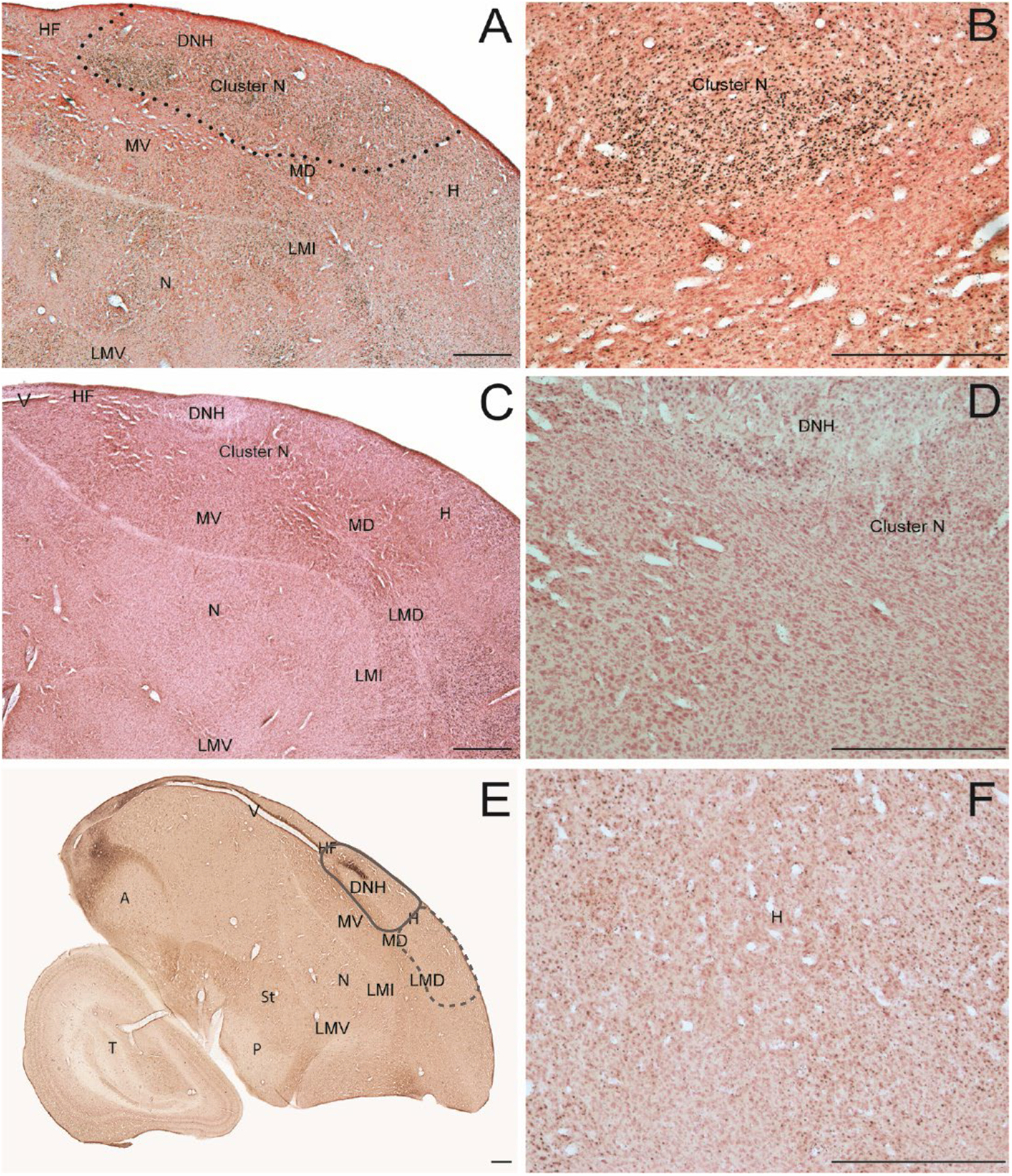
Right hemisphere sagittal sections of Swainson’s thrush. A) night-active bird with ZENK expression. Cluster N (dashed line) region indicated. (B) high magnification of cluster N region from (A). (C) day-active bird with ZENK expression. (D) high magnification of Cluster N region from (C). (E) GluR1 expression. (F) high magnification region of anterior hyperpallium used for control ZENK expression counting frame from (A). Abbreviations: A, acropallium; DNH, dorsal nucleus of the hyperpalllium; H, hyperpallium; LMD, lamina mesopallium dorsale; LMI, lamina mesopallium intermediate; lX, area X of the striatum; MD, dorsalmesopallium; MV, ventral mesopallium; N, nidopallium; P, pallidum; RA, robust nucleus of the acropallium; St, striatum; T, tectum; V, ventricle. Scale bars – 500μm.

### 3D reconstruction

We scanned stained serial brain sections using a Zeiss Axiocam 2 microscope system at 2.5x magnification and used Neurolucida software (MBFBioscience, Williston, VT) to create 3D reconstructions of regions of interest in the forebrain. To aid in visualization, outlines of the hyperpallium, mesopallium, ventricle, Cluster N, and DNH of each section were established. A 3D rendering was performed from these outlines from spaced sagittal sections through the right hemisphere of 2 birds. Cluster N was outlined inclusive of continuous regions of postive ZENK expressing cells in individual sections based, similar to previous investigations [22,26,31].

## Results

### Increased neural activity in the Cluster N region of night-active birds

We examined ZENK expression in the cortex of four night-active birds and two day-active birds. All night-active birds were exhibiting migratory restlessness immediately before anesthetization and perfusion. Fig 1A shows ZENK expression in a representative section for one night-active bird. Robust ZENK expression was generally observed broadly across large regions of the posterior hyperpallium (H), hippocampal formation, and dorsal mesopallium (MD).

Collectively, these regions form an area that was previously termed Cluster N [31]. This pattern of ZENK expression in Cluster N regions was consistent for all night-active birds. Fig 1B is a higher resolution image from 1A that illustrates the density of Cluster N cells expressing ZENK that tended to be grouped in clusters throughout the region. In contrast, there was little to no neural activity observed in day-active birds throughout Cluster N regions, as shown in Figs 1C. A higher resolution of the cluster N region from 1C that illustrates the paucity of ZENK expression in the day-active birds is shown in Fig 1D. The night vs day active neural activation was a defining feature of Cluster N boundaries and was further quantified as described below.

In addition to the ZENK expression in Cluster N, there was additional neural activation observed in the night-active birds in several cortical regions. The most prominent of these in the medial pallium included the posterior hippocampus, the posterior region of the nidopallium lying below the ventral mesopallium lamina, the entopallium, and the nucleus basalis. In the lateral pallium, neural activation was observed in the hippocampal formation, throughout the anterior pole of the hyperpallium, the ventral part of the frontal nidopallium, and the entopallium. These regions were also activated in the day-active birds.

In order to better understand the regional location of the ZENK expression being observed, adjacent brain sections were reacted for GluR1, as shown in Fig 1E. GluR1 expression indicates the location of AMPA glutamate receptors and has been well described by previous work in other bird species [10,19,39]. Generally, movement related neural activity is indicated by increased GluR1 expression. In the night-active birds we examined, GluR1 was highly expressed in the DNH but not in regions of the Cluster N.

The former qualitative observations of increased ZENK expression in the forebrain of night-active thrushes is supported by a quantitative analysis as well. We compared the relative ZENK expression using pixel density between night- and day-active birds for the Cluster N region and a separate region of the anterior hyperpallium and mesopallium that did not include Cluster N, as shown in Fig 1F (higher resolution from area in anterior hyperpallium from Fig 1A). We observed that the relative pixel density was significantly higher in night-active bird regions vs day-active birds (linear mixed model with bird included as a random variable, χ2 = 10.49, p-value = 0.0012), as shown in Fig 2. We next examined the relative expression between left- and right-hemispheres for ZENK activation. There was no difference in neural activation observed between the hemispheres (one sample t-test, sample estimate = 0.037, t = 1.84, p-value =0.16; Fig 2).

**Fig 2.**
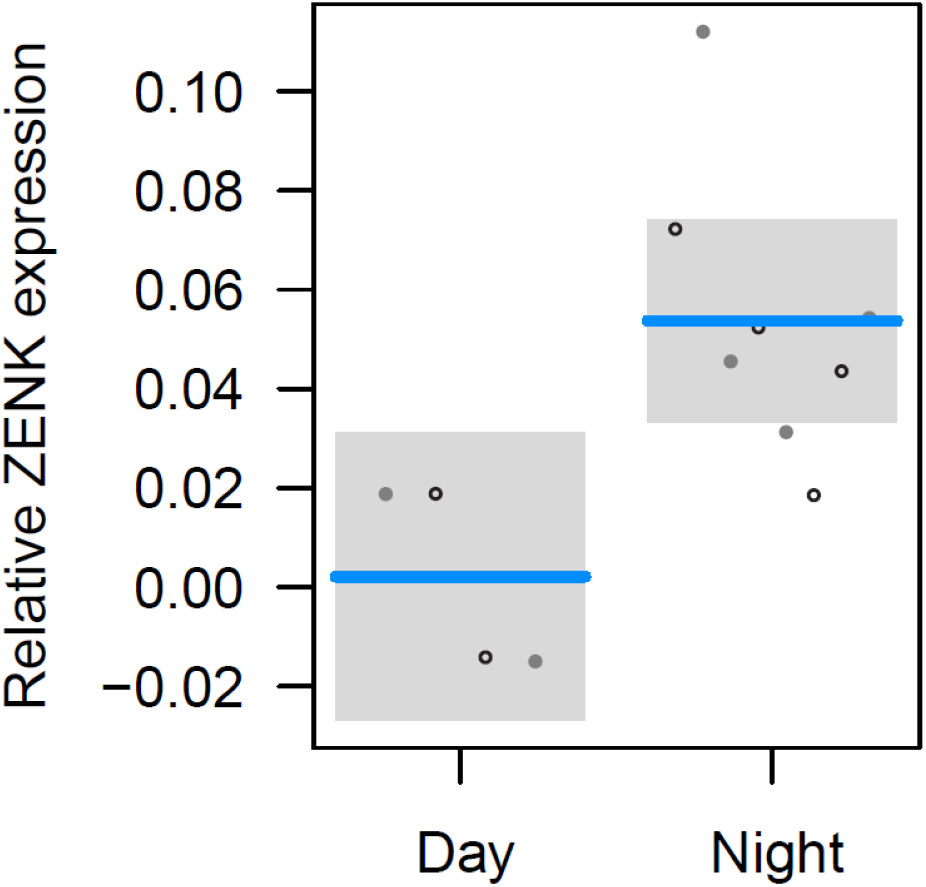
Relative ZENK expression in Cluster N for day- and night-active birds, showing significantly higher expression in night-active birds (** indicates significant difference between groups at p-value = 0.0012). Each point represents estimates for the right (open) and left (filled) hemispheres.

### 3D characterization of Cluster N

We next wished to determine the regional extent of Cluster N neural activation in the forebrain of the night-active birds during their migratory behavior patterns. We examined multiple alternate sections through the entire right hemisphere of two night-active birds, noting the regional boundaries of ZENK expression as well as the cytoarchitectural borders of different pallial regions. Boundaries of several regions in the pallium were traced using the Neurolucida 3D reconstruction software. As shown in Fig 3, Cluster N (dark green) encompassed large portions of the posterior hyperpallium and dorsal mesopallium in the thrush forebrain. More specifically, there was extensive neural activation observed in the areas of the hippocampal formation (HF) immediately anterior to the end of the lateral ventricle, surrounding the dorsal nucleus of the hyperpallium (DNH), and extending through the entire posterior hyperpallium. Activated neurons were observed from the surface of the pallium to extend ventrally into the upper regions of the dorsal mesopallium. No ZENK expression was observed in the ventral mesopallium (Fig 3., aqua). This pattern of neural ZENK expression was observed from near the midline to approximately 2.5 mm lateral to the midline (approximately halfway from medial to lateral in the forebrain) and occupied the majority of the posterior hyperpallium.

**Fig 3.**
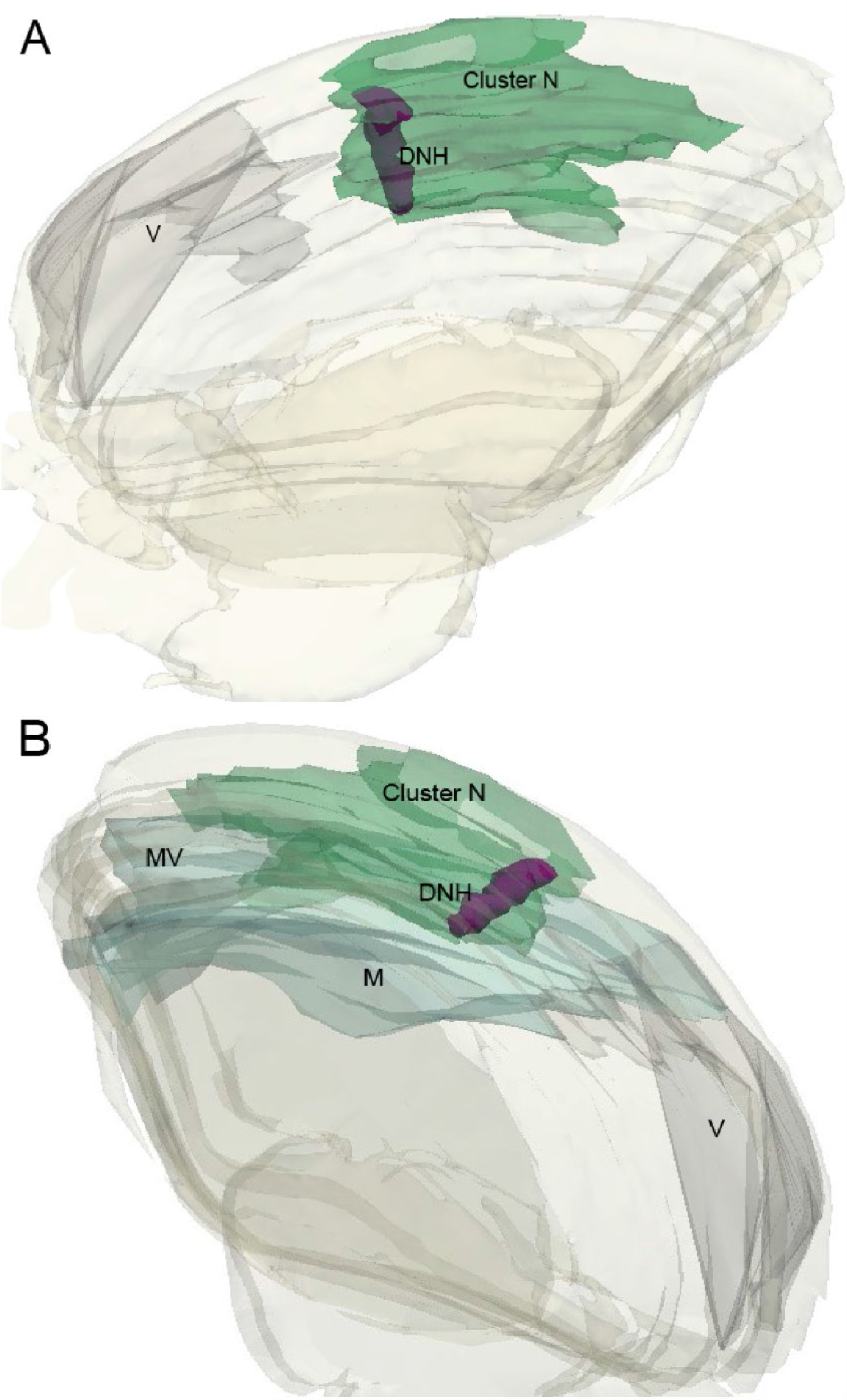
Right hemisphere 3D reconstructions of cortical regions containing Cluster N in a single night-active bird. (A) Medial to lateral sagittal reconstruction showing Cluster N (dark green) and the DNH relative to the ventricle. (B) Lateral to medial sagittal reconstruction showing Cluster N and the DNH (purple) relative to the mesopallium (aqua) and the ventricle (gray). Abbreviations: DNH, dorsal nucleus of the hyperpallium; M, mesopallium; MV, ventral mesopallium; V, ventricle.

## Discussion

Results from our study show that Swainson’s thrushes possess a cluster N region of the forebrain that is activated during night migratory behavior. Similar findings were recently reported for another North American bird, the white throated sparrow, as well as a number of European bird species [6,17,26,31,46]. Accordingly, this brain region appears to be highly conserved in birds, likely performing the same function across distinct migratory systems. Brodbeck et al. [6] found strong Cluster N activation only in night-birds engaged in migratory restlessness behavior and not in night birds that were at rest, suggesting that Cluster N is involved in navigation or orientation tasks and not just night vision [6]. We also found that the Cluster N region to be activated bilaterally, albeit our comparison sample size was small. These results of bilateral activation are consistent with those reported for sparrows and do not appear to support previous descriptions of lateralization specificity for night-activation of Cluster N in European robins [6,26]. We found that the Cluster N region of the forebrain included regions of the posterior hyperpallium, and dorsal mesopallium similar to that described for other bird species [18,26,31,46]. We also observed ZENK expression in the hippocampal formation of night-activated animals, similar to other findings in Garden warblers [23].

The regional location of Cluster N in these animals was extensive. It occupied nearly one third of the central dorsal forebrain, as revealed through our 3D reconstructions. Cluster N extended from the anterior end of the lateral ventricle to the end of the posterior hyperpallium in the anterior-posterior dimension. Extending from the midline, Cluster N lengthens anteriorly into more anterior regions of the hyperpallium, then recedes posteriorly through the lateral extent of the forebrain. Cluster N was also observed to fully surround the dorsal nucleus of the hyperpallium, similar to other species [22,31]. Although some cells within the DNH were activated in the night-active birds, the intensity of ZENK expression was noted to be much less than the surrounding Cluster N regions. The DNH appears to be a motor nucleus, since there was also noted to be intense GluR1 activation associated with high glutamate release (Fig 1E).

However, specific neural recording studies targeting DNH neurons have not been reported to the best of our knowledge, so the primary function of the DNH remains to be determined.

We did not document any evidence for lateralization of Cluster N in the Swainson’s thrush. Neural activity was elevated in both hemispheres of the brain in night-active birds, similar to the recent findings in North American sparrows [6]. It is possible that with a larger sample size we could detect lateralization but for now our results suggest that if it exists it is not very strong. These results suggest that cortical activation was not dependent upon unilateral visual or possibly magnetic receptor activation and is consistent with previous behavioral research. For example, Hein et al. [20,21] assayed the orientation of garden warblers and European robins (respectively) under both natural and changed magnetic fields when one eye was covered. Birds oriented correctly regardless of which eye was covered. Similar findings were obtained by Wilzeck et al. [43] using pigeons and Engels et al. [18] in European robins.

Our results do not support work conducted by Liedvogel et al. [26] with European robins. These authors examined ZENK expression while covering one or the other eye and showed that birds with their left eye covered had higher activity in cluster N than those with their right eye covered. They concluded that there was a specialized lateralization of function for cluster N in the right hemisphere. Future work with Swainson’s thrushes could employ a similar eye-blocking protocol, combining our assay for neuronal activity with an eye cover study.

What is the function of Cluster N? Mouritsen and colleagues [31] were the first to characterize the region as it relates specifically to migratory behavior. These authors defined the Cluster N region as being activated only at night in both European robins and garden warblers during their natural migration season, while the animals were sitting still. Similar conditions produced no activation in the same forebrain regions in two non-migratory birds, zebra finches and canaries [31]. Subsequent investigations have also identified a Cluster N region in additional European migratory birds during night migratory behavior [22,45,46]. Our work reported here, as well as another recent study, extend these findings to two North American migratory species, Swainson’s thrush and white throated sparrows, to show that Cluster N appears to be a conserved brain region involved in night migratory behavior generally for many birds [6].

Here our findings support previous evidence that links Cluster N activity to visual processing during the night. We observed a significant difference in ZENK expression in Cluster N between night and day-active birds, consistent with previous studies [6,31,45]. Further, when retinal signals were reduced by placing light-tight eye caps on European robins during migratory behavior, ZENK expression in Cluster N was significantly reduced [31]. As previously suggested, Cluster N appears to be a specialized region of the visual Wulst in birds [22,31]. As such, Cluster N would be included in the regions of the hyperpallium that incorporate the thalamofugal pathway that receive visual information from the retina through neurons in the optic thalamic nuclei [25,33]. Neural recordings from different bird species have identified responses in the posterior hyperpallium related to visual object recognition, visual motion, and visual disparity [1,4,7,16,35]. Many of these recordings appear to be located in corresponding Cluster N locations [5,7,35]. Yet, in all of this work, visual responses to night vs day luminence levels have not been compared. Still to the best of our knowledge, no studies have yet examined Cluster N neurons during night conditions with migratory species, so it is unknown if there is a specialization for low light responses from cells located in these regions.

It has also been suggested that Cluster N may be a specialized region for night navigation using both visual and geomagnetic informational cues [31,45,46]. We observed ZENK expression in the hippocampal formation in birds exhibiting night migratory restlessness, similar to other avian studies and the hippocampus has been repeatedly shown to be involved in spatial memory and navigation functions [23,24,38]. Zapka and colleagues (2009) lesioned Cluster N in migratory songbirds and disrupted magnetic orientation behavior but not sun or star compass behavior. Heyers et al. [22] used anatomical tracing to elucidate overlapping anterograde tracing from the retina to dorsal geniculate complex (Gld) in the thalamus with retrograde tracing from Cluster N in night migratory garden warblers. These authors further reported that some neurons in the of these neurons in Gld were disynaptic between retina and Cluster N; illustrating a potentially important thalamofugal pathway involved in night migratory behavior. Brodbeck et al. [6] found that Cluster N cells are activated at night, but only when birds are exhibiting migratory restlessness behavior and not when they are resting. In terms of Cluster N being involved in processing of geomagnetic information for navigation, only a few studies have reported indirect evidence to suggest a role. Heyers et al. [22] used ZENK expression during magnetic field manipulation to identify activated neurons in Cluster N in the same regions that received Gld projections. Wu and Dickman [44] manipulated both the direction and intensity of Earth strength artificial magnetic fields and C-fos expression to visualize a broad region of posterior hyperpallium neural activation in pigeons. Although Cluster N as a distinct forebrain region in pigeons has not yet been established, comparisons between hyperpallial regions activated by direct magnetic field stimulation in their pigeon study and those reported for night migratory birds appears similar [31,45,46]. Further, although a non-migratory species, pigeons have been shown to use Earth magnetic cues for natural homing behavior [15,30,40]. To date, no direct neurophysiological investigations examining Cluster N neurons to magnetic field manipulation have been reported, so it remains to be determined this intriguing brain region is involved in migratory behavior.

It was previously shown that magnetoreception is an important cue for migration in the Swainson’s thrush [8,9]. The present work represents the first step in elucidating the role of Cluster N in migratory navigation in these birds. The Swainson’s thrush includes two subspecies (coastal and inland). These subspecies hybridize in western North America but migrate along different routes, directly south to southern Mexico and central America or southeast, over the Rocky Mountains, to South America [11]. Pure and hybrid thrushes can be maintained in captivity and genomic tools (e.g., a highly contiguous reference genome) have already been developed for this species [12,14]. Accordingly, comparisons of transcriptomic data from day- and night-active birds from each subspecies and their hybrids could be used to identify genes that propagate signals between the retina, Cluster N and downstream targets that not only allow birds to orient on migration but also make different decision on which direction they will take.

Swainson’s thrushes are also large enough to carry some of the most advanced animal tracking technologies available today and direct tracking data exist for these birds already [11,13].

Accordingly, any candidates identified in the former captive work could be validated in free-flying birds.

